# Northstar enables automatic classification of known and novel cell types from tumor samples

**DOI:** 10.1101/820928

**Authors:** Fabio Zanini, Bojk A. Berghuis, Robert C. Jones, Benedetta Nicolis di Robilant, Rachel Yuan Nong, Jeffrey Norton, Michael F. Clarke, Stephen R. Quake

**Affiliations:** Department of Bioengineering, Stanford University, Stanford, CA, USA; Prince of Wales Clinical School, Lowy Cancer Research Centre, University of New South Wales, Sydney, Australia; Department of Oncology, Stanford School of Medicine, Stanford, CA, USA; Department of Immunology, Genetics and Pathology and SciLifeLab, Uppsala University, Uppsala, Sweden; Department of Surgery - Stanford Pancreas Cancer Research Group, General Surgery, Stanford University School of Medicine, Stanford, CA, USA; Department of Applied Physics, Stanford University, Stanford, CA, USA; Chan Zuckerberg Biohub, San Francisco, CA, USA

**Author notes:** these authors contributed equally (Fabio Zanini, Bojk A. Berghuis).

## Abstract

Single cell transcriptomics is revolutionising our understanding of tissue and disease heterogeneity, yet cell type identificationl remains a partially manual task. Published algorithms for automatic cell annotation are limited to known cell types and fail to capture novel populations, especially cancer cells. We developed northstar, a computational approach to classify thousands of cells based on published data within seconds while simultaneously identifying and highlighting new cell states such as malignancies. We tested northstar on human glioblastoma and melanoma and obtained high accuracy and robustness. We collected eleven pancreatic tumors and identified three shared and five private neoplastic cell populations, offering insight into the origins of neuroendocrine and exocrine tumors. northstar is a useful tool to assign known and novel cell type and states in the age of cell atlases.

## Introduction

The widespread adoption of single-cell transcriptomics has led to a growing body of “atlas” datasets describing tissue heterogeneity with unprecedented resolution [1]. As atlas data across species [2], tissues [3–7], development [8], and aging [4,9] are being amassed, a number of algorithms with or without deep learning have been proposed to leverage atlases for cell type classification [10,11]. The basic concept behind these efforts is to emulate the Basic Local Alignment Search Tool (BLAST) for nucleic acid and protein sequence matching [12], except in gene expression space.

Despite these recent approaches, by far the most common pipeline used by biomedicine laboratories to understand the composition of a single cell transcriptomic dataset is unsupervised clustering (e.g. Louvain [13] and Leiden [14] algorithms) as provided for instance by the 10X Genomics cellranger pipeline. After clustering, differential expression at the cluster level is used to identify marker genes and eventually annotate each cell population in the dataset via manual inspection.

A key reason why unsupervised clustering is still appealing is that a new dataset may contain novel cell populations. Because supervised (including deep) learning algorithms such as scVI rely heavily on training data and cannot cluster “outlier cells” on their own, the risk of misannotating the most interesting cells as a known cell type is real and bears profound consequences for the biological interpretation. This risk is particularly high when analysing tumor samples, where “novel” neoplastic cell states are not only possible but expected: the annotation of a malignant cluster as a healthy cell type is not acceptable.

Here we present northstar, an algorithm and software package that classifies single-cell transcriptomes guided by training data but is also able to discover new cell types or cell states. The inspiration for northstar comes from batch correction techniques [15,16], however northstar is much faster than those algorithms and its output is easier to interpret. To benchmark northstar we analyzed published datasets on glioblastoma [17] and melanoma [18] and found a superior ability to classify known and cluster novel cell types. We then applied northstar to newly collected 1,622 cells from 11 pancreatic cancer patients and rapidly classified them into healthy cell types and neoplastic states and ultimately gained useful biological insight into the composition and origin of the tumors.

## Results

### northstar identifies cell types guided by a cell atlas

northstar is a computational approach to identify cell types in a new single cell dataset leveraging one or multiple cell atlases. The unique feature of northstar is that every new cell can be either assigned to either a known atlas cell type or a novel cluster. An implementation in C++/Python is available at https://github.com/northstaratlas/northstar and preprocessed atlases for immediate use are available at https://northstaratlas.github.io/atlas_landmarks.

As input, the gene expression table for the new dataset and the gene expression table and cell type annotations for the cell atlas are specified by the user. northstar can use either an average expression vector for each cell type or a small subsample of each cell type. Atlas averages and subsamples can be used by northstar as *reference landmarks* to annotate the new cells. In this sense, northstar serves the same purpose in single-cell datasets as the North Star always had for maritime navigation: providing fixed points that guide the exploration of new landscapes. To simplify adoption, we provide precomputed landmarks (averages and subsamples) of several atlases (see above link). If a precomputed atlas is chosen, the user only needs to specify its name: counts and annotations are downloaded automatically.

The algorithm consists of the following steps. First, atlas landmarks (averages or subsamples) are merged with the new single-cell dataset into a single data table (Figure 1A). Then, informative genes are selected: upregulated markers of each atlas cell type are included as well as genes showing a high variation within the new dataset. A similarity graph of the merged dataset is constructed, in which each edge connects either two cells with similar expression from the new dataset or a new cell with an atlas cell type (Figure 1B). Finally, nodes in the graph are clustered into communities using a variant of the Leiden algorithm that prevents the atlas nodes from merging or splitting [14]. The output of northstar is an assignment of each cell to either an atlas cell type or, if a group of cells show a distinctive gene expression profile, to a novel cluster (Figure 1C).

**Figure 1.**
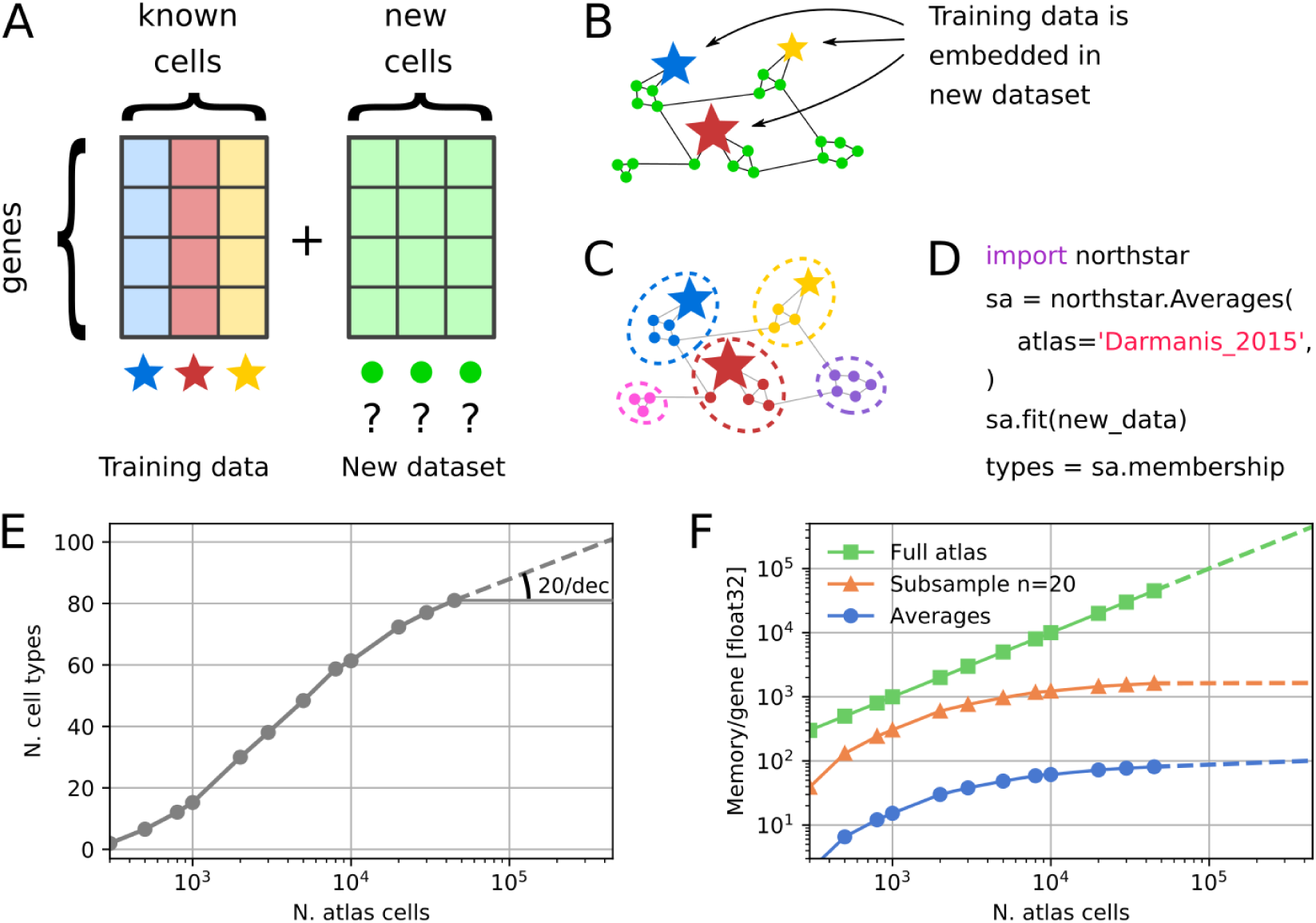
northstar concept and scalability. **(A)** northstar’s input: the gene expression table of the tumor dataset and the cell atlas. Annotated cell type averages are depicted by colored stars, unannotated new cells by green circles. **(B)** Similarity graph between atlas and new dataset. **(C)** Clustering the graph assigns cells to known (stars) or new cell types (pink and purple and circles, bottom left and right). Cell types themselves do not split or merge. **(D)** Typical code used to run northstar. **(E)** Number of cell types with at least 20 cells in the Tabula Muris atlas (FACS data only), subsampled to different number of cells [2]. **(F)** Memory requirements for northstar barely increase with larger training atlases.

northstar is designed to be easy to use (Figure 1D). To examine northstar’s scalability to large atlases, we downloaded the Tabula Muris plate data [2], subsampled it to different cell numbers, and counted the number of cell types with at least 20 cells. As more cells were sampled new cell types were discovered, however with diminishing returns. At full sampling (50,000 cells), we estimated at most 20 new cell types per tenfold increase in cell numbers (Figure 1E). Because of this sublinear behaviour, northstar scales to atlases of arbitrary size, unlike a naive approach that combines all atlas cells with the new dataset (Figure 1F). Although subsampling each cell type (e.g. 20 cells) requires more storage memory than a single average, their scaling behaviour is exactly the same (i.e. logarithmic or better). In practical tests, we found that subsampling helps with very heterogeneous cell types, however it is more reliant on a high-quality atlas annotation.

### Benchmark against published datasets on healthy brain and glioblastoma

To validate northstar’s performance, we analyzed a glioblastoma (GBM) dataset [17] on the basis of a previously annotated cell atlas of the human brain by the same authors [3]. The annotations of the GBM dataset according to the original authors define seven healthy cell types: neurons, astrocytes, oligodendrocytes, oligodendrocyte progenitor cells (OPCs), endothelial cells, microglia, and other immune cells. In addition to these seven, an additional cluster of neoplastic cells is described. Of these cell types, the first five are also present in the brain atlas, while some fetal cells were excluded from the atlas because the GBM patients were adults. A projection of the GBM data via t-Distributed Stochastic Neighbor Embedding (t-SNE) is shown in Figure. 2A with the original annotations. The relative abundances of the various cell types in the atlas and GBM data is shown in Figure 2B.

**Figure 2.**
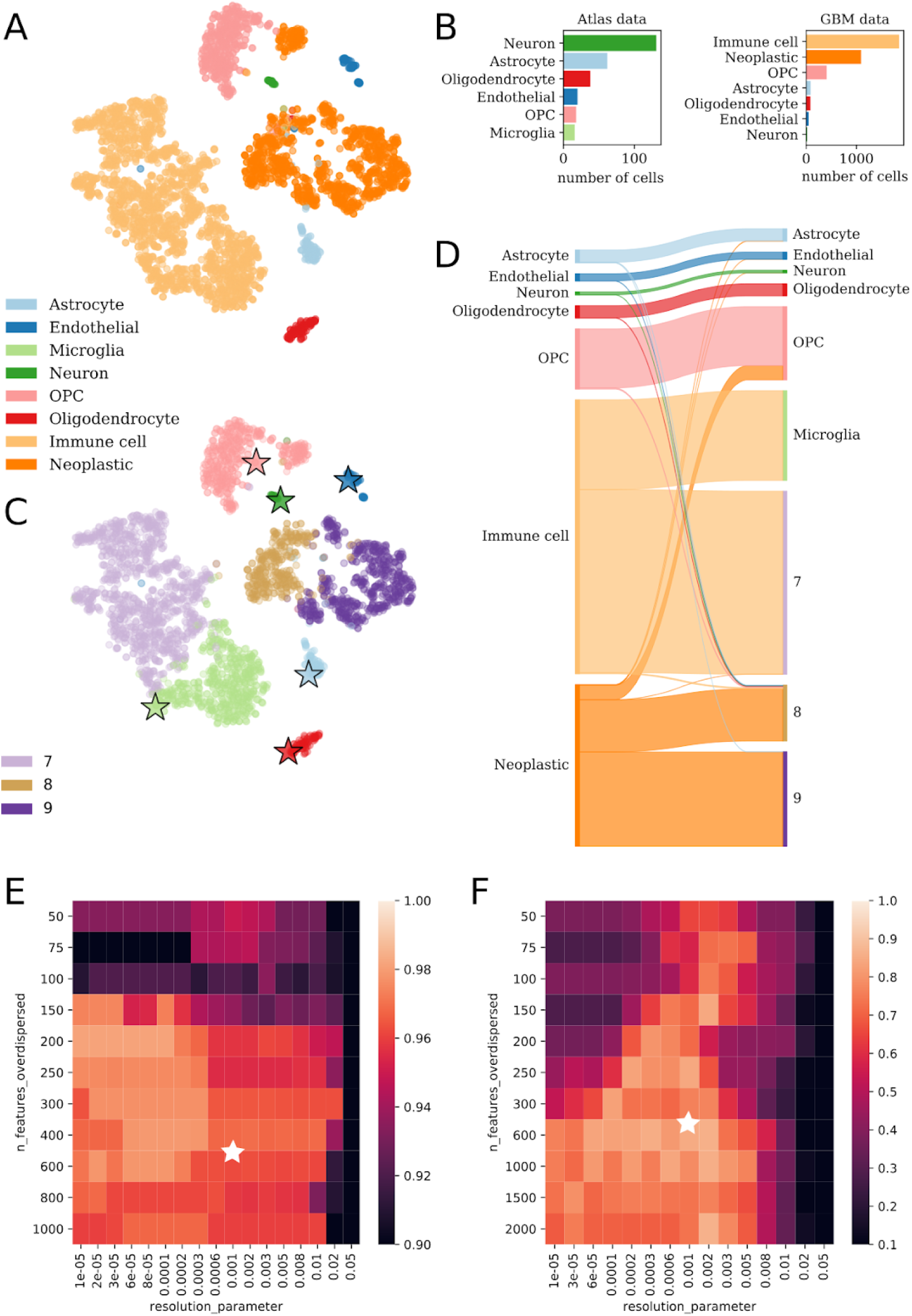
Annotation of glioblastoma and melanoma. **(A)** t-SNE of all cells in Darmanis et al. (2017) colored by original annotation [17]. **(B)** The number and type of cells that constitute the atlas data and the new glioblastoma data. **(C)** The same embedding of glioblastoma cells as in (A) but colored by northstar’s annotation and showing atlas cells (stars: atlas averages). **(D)** Sankey diagram with original manual annotations (left) vs. automated annotation by northstar (right). Band widths are proportional to the number of cells in the cluster. (E-F) Fraction of correctly assigned cells to known cell types (E) in the GBM dataset [17] (mean of 3 northstar runs) and (F) in an independent study on melanoma in mice [18] (mean of 5 runs) upon variation of the number of overdispersed features from the new dataset and the resolution parameter of the Leiden clustering. White stars indicate default parameters.

We calculated cell type gene expression averages of the brain atlas, deleted the annotations from the GBM data, and fed labeled atlas averages and unlabeled GBM data into northstar. In less than 9 seconds of runtime on a modern laptop, we obtained a classification of the new cells and observed that our method recapitulated previously reported cell types and also created new classes for myeloid and neoplastic cells as these cell types were absent from the atlas (Figure 2C, Supplementary Figure 1).

A detailed analysis of the original annotations versus new annotations generated by our algorithm highlights the strength of the method (Figure 2D). Almost all cells of known types were correctly assigned to their respective cluster. The main misclassification was a population annotated as neoplastic in the original paper but classified as oligodendrocyte precursor (OPC) by northstar. However, those cells formed a relatively distinct cloud in both Figure 2C and, even more clearly, in the original t-SNE plot (Supplementary Figure 1). Moreover, a diverse population of immune cells is either classified as microglia, which are resident immune cells, or into new cluster 7. Neoplastic cells were correctly assigned to new clusters 8 and 9. To identify why both immune and neoplastic clusters were split by northstar we examined their connectedness in the similarity graph. We found that their subcomponents were only weakly connected (Supplementary Figure 2); the Leiden algorithm - which wasn’t available to Darmanis et al. in 2017 - split them to increase internal connectedness [14]. Differential expression analysis indicates that the immune clusters express CD207, CD300E and other classical immune surface markers while the neoplastic cell clusters expressed KLK6, MIR1322, and other RNAs previously implicated in malignancies [19,20].

To test northstar’s accuracy and stability, we repeatedly classified the glioblastoma dataset in 10 independent runs with default parameters and found that in average 97% of cells were correctly classified with a standard deviation between runs smaller than 1%, indicating both high accuracy and stability. Similar classifications were obtained using northstar with a subsample of the atlas, confirming that the classification is robust (Supplementary Figure 3). To test northstar’s sensitivity to the input parameters, we performed thousands of runs - using an atlas subsample - covering a wide range of the following parameters: (1) number of marker genes per cell types, (2) number of overdispersed features from the new dataset, (3) number of principal components retained, (4) number of graph neighbors, and (5) resolution parameter of the Leiden clustering. We found that only (2) and (5) had any detectable impact on the fraction of correctly assigned cell types, and that our classification was very accurate as long as the number of overdispersed features from the new dataset is at least 150 (to enable sufficient discrimination) and the resolution parameter is less than 0.01 (to avoid subcluster splitting) (Figure 2E).

**Figure 3.**
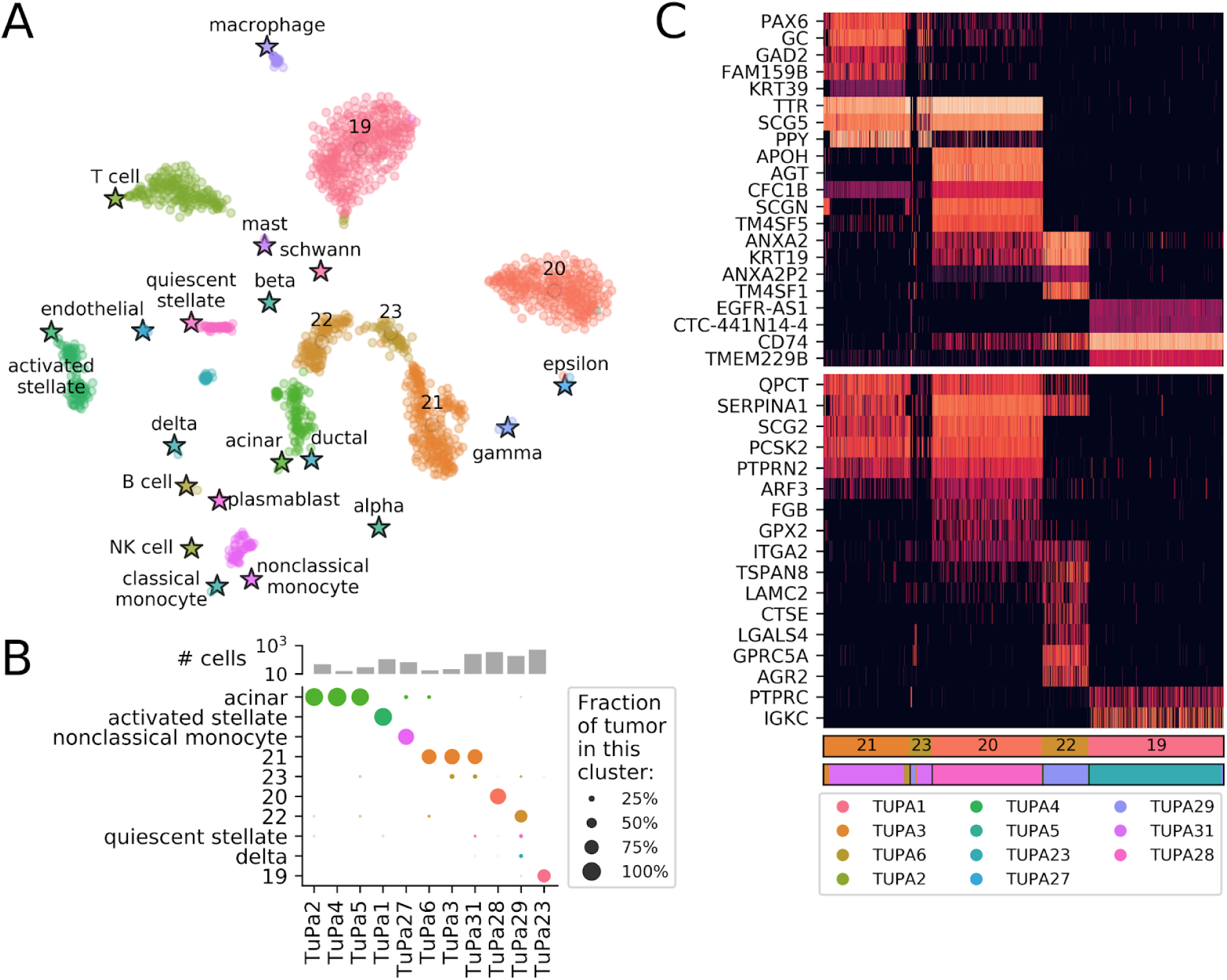
Annotation of new pancreatic tumor samples. **(A)** t-SNE of 11 pancreatic tumors together with averages from two atlases by Baron et al (2016) and Zanini et al. (2018) [22,23]. Stars: atlas averages. **(B)** Number of cells from each tumor and fractions of cells belonging to each cell type. **(C)** Top differentially expressed genes for each novel cluster (top) and some known markers (bottom) for pancreatic cancers [35]. Black indicates no expression, red to white increasing expression levels. PTPRC (CD45) and IGKC are added to highlight the probable B cell phenotype of cluster 19. First bar: novel cluster (colored as in A and B). Second bar: patient of origin, with legend below.

To test northstar’s performance on a larger dataset, we analyzed 4,626 cells from a melanoma murine model focused on infiltrating immune cells [18]. As for the GBM data, we found high accuracy with the default parameters (~90% correctly assigned immune cells using Tabula Muris marrow annotations only). For this dataset, selecting 600 or more features from the new dataset generally increased accuracy (Figure 2F). In terms of speed, northstar classified the >4,000 cell melanoma dataset in approximately 9 seconds on a modern laptop using a single low-voltage CPU core (Supplementary Figure 4). These findings highlight northstar’s core concept that even just a few example transcriptomes for each cell type are sufficient to guide the classification task robustly and efficiently.

### Classification of healthy and neoplastic cells in pancreatic cancer

To test the ability of northstar to provide biological insight into new datasets, we collected 1,622 single cells from 11 pancreatic tumors (Supplementary Table 1) and aimed to determine their composition in terms of known cell types and, potentially, novel clusters. Briefly, tumors were surgically resected at Stanford hospital from patients with diagnosed pancreatic cancer and dissociated into single-cell suspensions (see <u>Methods</u>). Single cells were isolated by fluorescence-activated cell sorting (FACS) into 96- or 384-well microtiter plates and processed as described previously [4,21].

northstar was then used to identify the cells in these samples based on the pancreas atlas by Baron et al. (2016) [22] and the immune atlas by Zanini et al. (2018) [23]. We found that the cells from the cancer patients were classified into both known and novel cell types (Figure 3A). Among the known cell types were mesenchymal, exocrine and endocrine cell types, and various immune cells. Although all tumors were resected in a similar procedure, the fraction of cells belonging to each known cell type varied considerably among patients (Figure 3B and Supplementary Table 2). This reflects the challenges of isolating tumor tissue during surgery without capturing surrounding tissue, and making single-cell suspensions from pancreatic exocrine tissue while the tissue is self-digesting.

Samples TuPa2, 4, and 5 were composed mostly of acinar cells - or cells that closely resemble that cell type. In fact, it is clear from the embedding (Fig 3A) that many of these cells are close to both acinar and ductal types. This was expected for patients TuPa2 and 4 because the tumor was diagnosed as Pancreatic Ductal Adenocarcinoma (PDAC) but not TuPa5, which is annotated as a neuroendocrine tumor. TuPa1 was clinically described as fibromatosis and we found it is essentially composed of activated stellate cells. This is consistent with recent literature [24–26]. Sample TuPa27 was classified as composed mainly of monocytes, which was surprising considering that it was clinically described as a neuroendocrine tumor. Because surgery took longer than usual for this sample, we speculate that the tumor cells might have degraded leaving only the more robust immune cells. All other samples classified in one or more novel clusters. TuPa6, 3, and 31 shared cluster 21 though those patients were diagnosed with three different conditions: ampullary adenocarcinoma, mucinous cystic neoplasm, and neuroendocrine tumor, respectively. The embedding confirms northstar’s prediction and places these cells somewhere between alpha and gamma/PP endocrine types, indicating an endocrine origin. A minority of cells in samples TuPa3 and 31 belonged to another new cluster, 23, which is located nearby on the embedding. The tumor from TuPa28 belonged to its own private cluster 20. Its location is indicative of an endocrine lineage, in agreement with the patient’s diagnosis of neuroendocrine tumor. The donor of TuPa29 was the only one diagnosed with invasive adenosquamous carcinoma and was found to contain a majority of cells in a new, almost private cluster 22 located in the vicinity of acinar cells. A minority of cells from this sample were assigned to quiescent stellate and delta cells. Finally, sample TuPa23 was composed entirely of quasi-private cluster 19, which is surrounded in the embedding by resident immune cells.

To better understand the nature of the novel clusters 19 to 23, we computed differentially expressed genes (DEGs) between each novel cluster and all other cells by Kolmogorov-Smirnov statistics on the expression. In short, we searched for genes that are expressed by a high fraction of the cells within the focal cluster and by few cells outside of it (Figure 3C, top heatmap and Supplementary Table 3). The expression of PPY by clusters 21 and 23 is suggestive that these might be neoplastic cells derived from gamma/PP precursors. Clusters 21, 23, and 20 all express TTR and SCG5 but the latter cluster is missing PPY; this favors a distinct endocrine origin for the latter cluster. Cluster 22 expresses KRT19, indicating an epithelial origin which is consistent with their proximity on the t-SNE with acinar and ductal cells. Finally, cluster 19 expresses CD74 which is part of the MHC class II machinery found in immune, antigen presenting cells. To further validate these results, we also looked at the expression of known markers for endocrine and exocrine cancers (bottom heatmap) and found that clusters 20, 21 and 23 show an expression consistent with endocrine cancer cells, while cluster 22 is consistent with exocrine cancer. Cluster 19 expresses PTPRC and IGKC, indicating it is related to B cells. Its location on the embedding supports an immune cell type, although the atlas B cells and plasmablasts are not located in proximity. This discrepancy might be due by biological differences between tumor infiltrating B cells and peripheral blood lymphocytes from healthy subjects.

## Discussion

Annotating a new single-cell transcriptomic dataset traditionally involves clustering without an atlas, however this process is laborious and can be inaccurate. Geometric subsampling of the atlas [27], followed by merging with the new data and unsupersived clustering is a viable route, however known cell types can split into subclusters or merge into superclusters, leading to difficulties in interpretation. In our experience such cases happen often because clustering can be performed at different resolutions leading to equally valid classifications (e.g. all immune cells, lymphocytes, T cells). northstar improves over these approaches by combining cell-type aware subsampling of the atlas with fixed clustering resolution and a stable clustering algorithm that guarantees internal connectedness [14]. Cell types can neither split nor merge simply because they are determined by the atlas.

northstar is very efficient because it approximates a cell atlas by compressed representations, i.e. averages or small subsamples. One can easily use an atlas with millions of cells on a laptop with 16 GB of RAM as long as the number of ***cell types*** remains within a few thousands. Current atlases only have tens (see Figs. 2 and 3) or hundreds of cell types and that figure is unlikely to be surpassed by orders of magnitude. Classifying thousands of glioblastoma and pancreatic tumor cells only took seconds and is actually faster than computing their t-SNEs.

northstar clearly separates the concerns of building the similarity graph (see Figure 1B) and classifying cell types based on an existing graph (Figure 1C). Although a simple and effective algorithm for graph building is implemented, we intentionally allow the use of custom similarity graphs to cope with batch effects. A plethora of batch correction methods has been proposed and northstar is designed to be compatible with most of them [15,16,28,29].

Unfortunately, many cell atlases and single cell datasets are poorly disseminated. Data access is idiosyncratic to each dataset and often requires manual steps (e.g. writing emails to the authors). To change this trend and help disseminate northstar we provide a website with averages and subsamples for several atlases that can be accessed programmatically: https://northstaratlas.github.io/atlas_landmarks. This makes it easy to combine atlases and cherry pick different cell types from each to maximize the leverage provided by the annotations.

The analyses of brain and pancreatic tumors presented here highlights the utility of northstar to quickly characterize the cell type composition of tumors. Simple differential expression can be applied immediately afterwards (Figure 3C) to identify the nature of the new clusters and to shed light on their biological origins. Sampling human tumors is challenging due to cell death preferential to certain cell types (e.g. neurons, pancreatic exocrine cells), hence northstar is an ideal tool to verify whether the cell types of interest are captured effectively. Moreover, the joint analysis of multiple patient samples is informative about how stereotypic neoplastic cell state progression is across individuals. In the 11 pancreatic tumors analyzed, we observed both shared and private clusters and also found corroborating evidence linking fibromatosis to activated stellate cells [24–26]. Cell atlases from large numbers of cancers are being collected in addition to healthy tissues [30]. This will further boost the utility of northstar for rapidly classifying tumors into known or novel neoplastic cell states.

Cell atlases provide an invaluable resource to study heterogeneous disease and in particular cancer. northstar’s unique ability to identify healthy and neoplastic cells is an important step towards personalized diagnosis and characterization of disease states at the single cell level.

## Methods

All methods were carried out in accordance with relevant guidelines and regulations.

### Brain atlas, glioblastoma dataset, and pancreas atlas

Gene expression count tables and cell type annotations were downloaded from NCBI’s Gene Expression Omnibus website (brain atlas: <u>GSE67835</u>, glioblastoma: <u>GSE84465</u>, pancreas atlas: <u>GSE81547</u>). To combine the brain atlas and glioblastoma dataset, only genes that were present in both datasets were retained. Gene expression counts were normalized by 1 million total counts per cell. For the brain and pancreas atlases, arithmetic averages of the normalized counts were computed for each cell type and stored. Fetal cell types were excluded from the brain atlas since the glioblastoma dataset refers to adult patients, while ambiguous cell types (e.g. “unknown”, “hybrid”) were excluded from both atlases.

### Pancreatic cancer dataset

A novel single-cell transcriptomic dataset from human individuals with pancreatic cancer was collected. Eleven individuals were sampled with a total of 1,622 cells. A table of individual metadata is available as Supplementary Table 1.

Pancreatic tumor tissue was obtained at the Stanford Hospital from individuals undergoing surgery for pancreatic cancer between September 2015 and June 2018. Informed consent was obtained from all subjects/participants for the study. The study was approved by the Stanford Hospital Institutional Review Board for research on human subjects (IRB 4344). Single-cell suspension was then achieved by dissociating the samples for 1-2 hours with Collagenase/Hyaluronidase (Stemcell Technologies, 7912) in DMEM 1% FBS, followed by 2 minutes digestion with Trypsin-EDTA (0.25%, Life Tech, except for TuPa#1, #3 and #4). ACK and DNAse treatments were performed as needed.

Single cells were isolated by fluorescence-activated cell sorting (FACS) on FACS Aria II (BD Biosciences for TuPa#1-6) into a single well of 96-well plate and on Sony SH800Z (for TuPa 23,27,28,29,31) into single wells of 384-well Biorad HardShell plates. Antibodies used for sorting pancreatic cells were: EpCAM fluorescein isothiocyanate (FITC), CD49f phycoerythrin (PE), CD24 PE-Cy7, CD44 Allophycocyanin (APC), hCD45/GPA Pacific-blue (BioLegend). Cells were gated on the basis of forward- and side-scatter profiles, and live/dead discrimination was obtained with Sytox Blue (ThermoFisher #oS34857) or DAPI (4′,6-diamidino-2-phenylindole). The plates were pre-filled with 500nl of lysis buffer containing poly-T capture oligos, spike-in External RNA Controls Consortium (ERCC) control RNAs, and other molecules as described elsewhere [2]. cDNA synthesis and amplification was performed as described in the same reference. Libraries using in-house Tn5 were done as described and sequenced on Illumina NovaSeq 6000 on S2 or S4 flow cells and 100 base paired-end kits at an average depth of 1 million reads per cell. To avoid index hopping, dual unique barcodes with a reciprocal Hamming distance > 2 were used.

The sequencing read pairs were mapped to the human genome using STAR RNA aligner [31] and sorted by name. Genes were counted using HTSeq [32]. One of us (FZ) is the maintainer of HTSeq. A new package for grouping mapping and counting was developed by one of us (FZ) and is available on GitHub: https://github.com/iosonofabio/bag_of_stars. Cells with less than 100,000 reads were discarded.

### Data processing

For both datasets, count tables were further analyzed in Python 3.7 using numpy [33], pandas [34], and singlet. The latter package was developed by one of us (FZ) and is available on GitHub: https://github.com/iosonofabio/singlet and on PyPI: https://pypi.org/project/singlet/. A detailed description of the northstar algorithm is available in <u>Supplementary Text 1</u>.

## Acknowledgements

The authors thank Sai Saroja Kolluru, Norma Neff, Brian Yu, and staff on the Chan Zuckerberg Biohub sequencing team for assistance with library making and sequencing, Vincent Traag for help with Leiden, and all members of the Quake lab for helpful discussions. We also thank J Lyon and JE Manning Fox for processing pancreas samples at the Alberta Diabetes Institute IsletCore, Human Organ Procurement and Exchange (HOPE) and Trillium Gift of Life Network (TGLN) for the coordination of donor organs, and organ donors and their families for their generous contribution to research. This study was supported in part by the California Institute for Regenerative Medicine (CIRM) through the CIRM Center of Excellence in Stem Cell Genomics grant #GC1R-06673-A and the Chan Zuckerberg Biohub. B.A.B. was supported by a Rubicon Fellowship from the Netherlands Organization for Scientific Research. R. Y. N. was supported by an International Post Doc Grant from the Swedish Research Council.

## Data availability

northstar is available on GitHub at https://github.com/northstaratlas/northstar. Cell type averages and subsamples for a number of cell atlases are available at https://northstaratlas.github.io/atlas_landmarks/. All scripts used to generate the figures and tables for this manuscript are available on GitHub at https://github.com/northstaratlas/northstar_analysis/. The code is written in Python 3, C, and C++ and is tested via continuous integration on Linux and OSX.

## Supplementary Materials

**Supplementary Table 1.**
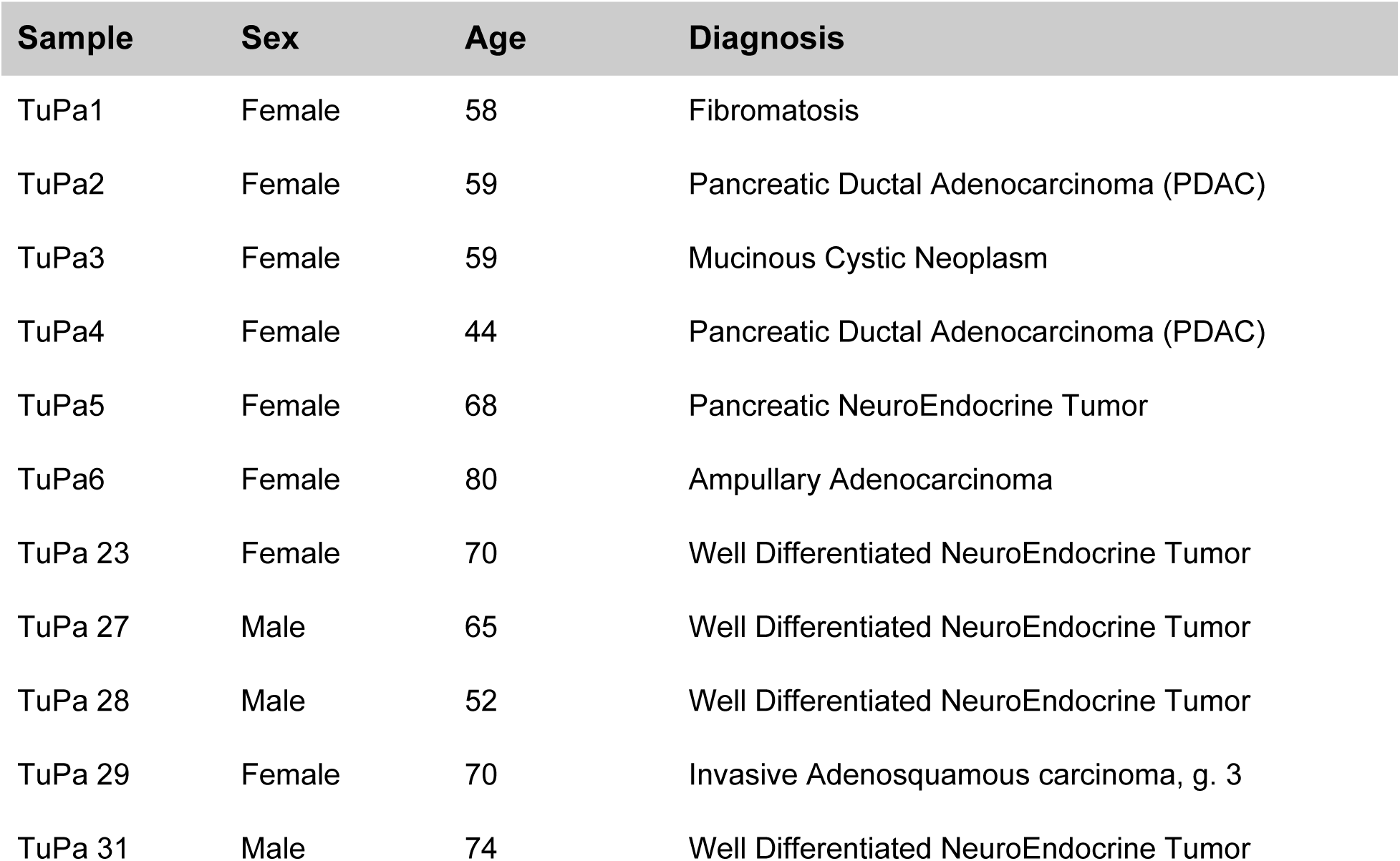
Patient metadata for pancreatic cancer dataset

| Sample | Sex | Age | Diagnosis |
| --- | --- | --- | --- |
| TuPa1 | Female | 58 | Fibromatosis |
| TuPa2 | Female | 59 | Pancreatic Ductal Adenocarcinoma (PDAC) |
| TuPa3 | Female | 59 | Mucinous Cystic Neoplasm |
| TuPa4 | Female | 44 | Pancreatic Ductal Adenocarcinoma (PDAC) |
| TuPa5 | Female | 68 | Pancreatic NeuroEndocrine Tumor |
| TuPa6 | Female | 80 | Ampullary Adenocarcinoma |
| TuPa 23 | Female | 70 | Well Differentiated NeuroEndocrine Tumor |
| TuPa 27 | Male | 65 | Well Differentiated NeuroEndocrine Tumor |
| TuPa 28 | Male | 52 | Well Differentiated NeuroEndocrine Tumor |
| TuPa 29 | Female | 70 | Invasive Adenosquamous carcinoma, g. 3 |
| TuPa 31 | Male | 74 | Well Differentiated NeuroEndocrine Tumor |

**Supplementary Table 2.** Number of cells from each pancreatic tumor sample assigned to each known cell type or new cluster.

| Cell type | TuPa |  |  |  |  |  |  |  |  |  |  |
| --- | --- | --- | --- | --- | --- | --- | --- | --- | --- | --- | --- |
|  | 1 | 2 | 3 | 4 | 5 | 6 | 23 | 27 | 28 | 29 | 31 |
| Macrophage |  |  |  |  |  |  | 3 |  | 19 |  |  |
| Schwann cell |  |  |  |  |  |  |  |  |  |  |  |
| Mast cell |  |  |  |  |  |  | 1 |  |  | 1 |  |
| T cell |  |  |  |  |  |  | 154 |  | 23 | 3 | 1 |
| NK cell |  |  |  |  |  |  |  |  |  |  |  |
| B cell |  |  |  |  |  |  | 1 |  |  |  |  |
| Plasmablast |  |  |  |  |  |  | 1 |  |  |  |  |
| Classical monocyte |  |  |  |  |  |  |  | 1 |  |  |  |
| Nonclassical monocyte |  |  |  |  |  |  |  | 58 |  |  |  |
| Endothelial | 3 |  |  |  |  |  |  |  |  |  |  |
| Quiescent stellate | 2 | 1 |  |  |  |  |  |  | 3 | 22 | 15 |
| Activated stellate | 105 |  |  |  |  |  |  |  |  |  |  |
| Ductal |  |  |  |  |  |  |  |  |  |  |  |
| Acinar |  | 46 |  | 16 | 28 | 2 |  | 9 |  | 3 |  |
| Alpha |  |  |  |  |  |  |  |  | 1 |  |  |
| Beta |  |  |  |  |  |  |  |  |  |  |  |
| Gamma/PP |  |  |  |  |  | 1 |  |  | 1 |  |  |
| Delta |  |  |  |  |  |  |  |  | 1 | 24 | 2 |
| Epsilon |  |  |  |  |  |  |  |  | 1 |  |  |
| 19 |  |  |  |  |  |  | 356 |  |  |  |  |
| 20 |  |  |  |  |  |  |  | 295 |  |  |  |
| 21 |  |  | 18 |  |  | 14 |  |  |  |  | 201 |
| 22 |  | 1 |  |  | 1 | 1 |  |  |  | 121 |  |
| 23 |  |  | 4 |  | 1 |  | 4 |  | 2 | 10 | 38 |

**Supplementary Table 3.** Top 15 differentially expressed genes per new cluster. Genes to which we could assign a clear association with known cell types are highlighted in boldface.

| Cluster |  |  |  |  |
| --- | --- | --- | --- | --- |
| 19 | 20 | 21 | 22 | 23 |
| EGFR-AS1 | APOH | PAX6 | **ANXA2** | **TTR** |
| CTC-441N14-4 | AGT | GC | C19orf33 | GC |
| C19orf57 | CFC1B | GAD2 | **KRT19** | SCG5 |
| **CD74** | SCGN | FAM159B | ANXA2P2 | **PPY** |
| TMEM229B | TM4SF5 | **KRT39** | TM4SF1 | RPL34P18 |
| **HLA-DRA** | CFC1 | **PPY** | OCIAD2 | **CHGB** |
| DNAJC27-AS1 | CALY | ERO1B | TMPRSS4 | PAX6 |
| PCDH10 | PCSK1N | **CHGA** | LGALS3 | TM4SF4 |
| TNR | CCL15 | SCG5 | PLAT | **CHGA** |
| GAREM2 | MGST1 | **TTR** | GPRC5A | ATP5EP2 |
| RP11-253M7-1 | SLC14A1 | **CHGB** | S100P | C10orf10 |
| **HLA-DPB1** | RTBDN | PAPPA2 | RPL7P1 | HSP90AA2P |
| **HLA-DRB1** | SPTSSB | CPE | RPL7P23 | ERO1B |
| ST7L | TPPP3 | SCG3 | RPL7P32 | PDK4 |
| **HLA-DRB5** | **COL28A1** | PDK4 | RPL7P6 | TTI2 |

**Supplementary Figure 1.**
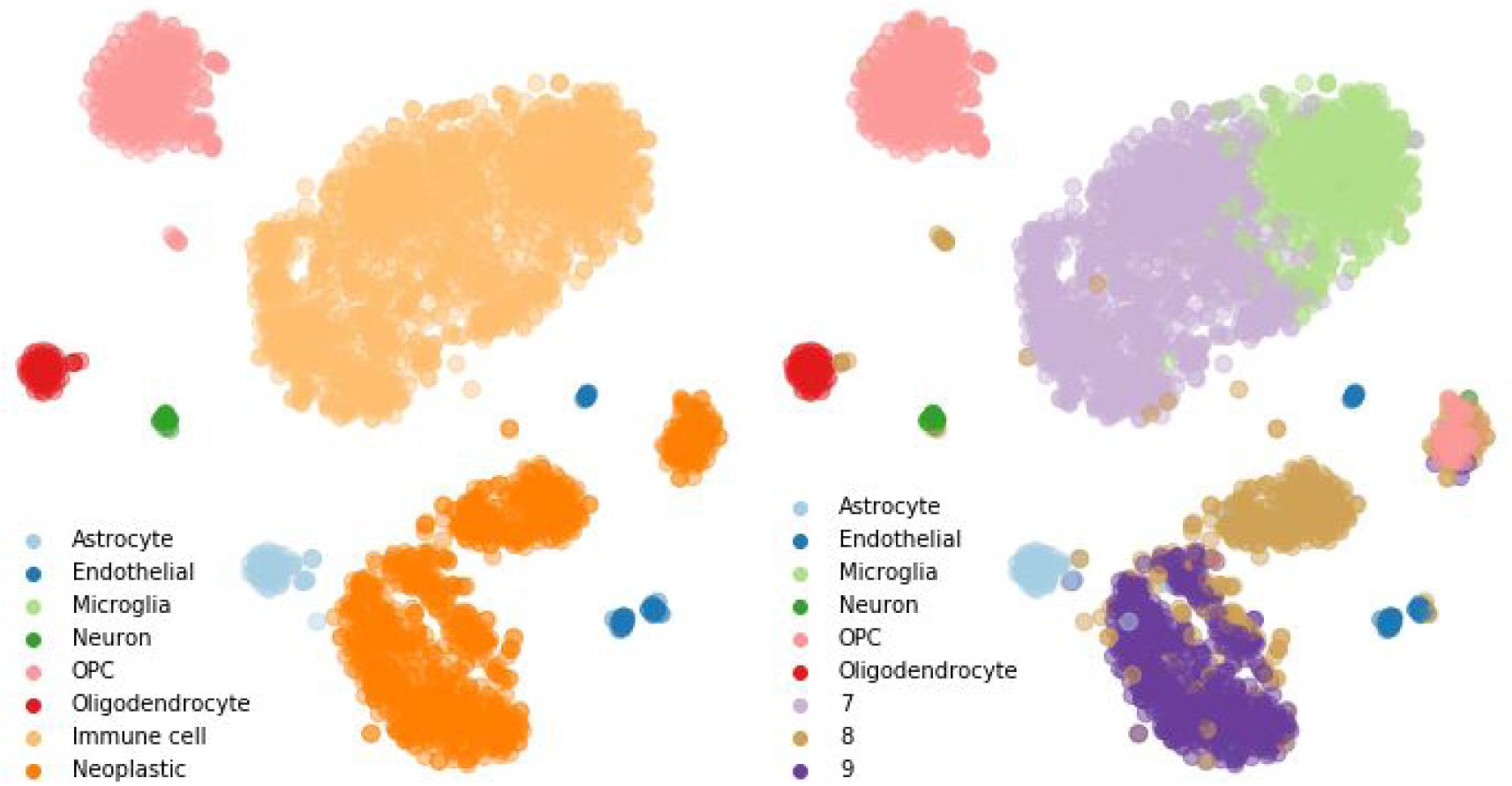
Original t-SNE representation of glioblastoma dataset. Cells colored by original annotation (left) and newly annotated classes (right)

**Supplementary Figure 2.**
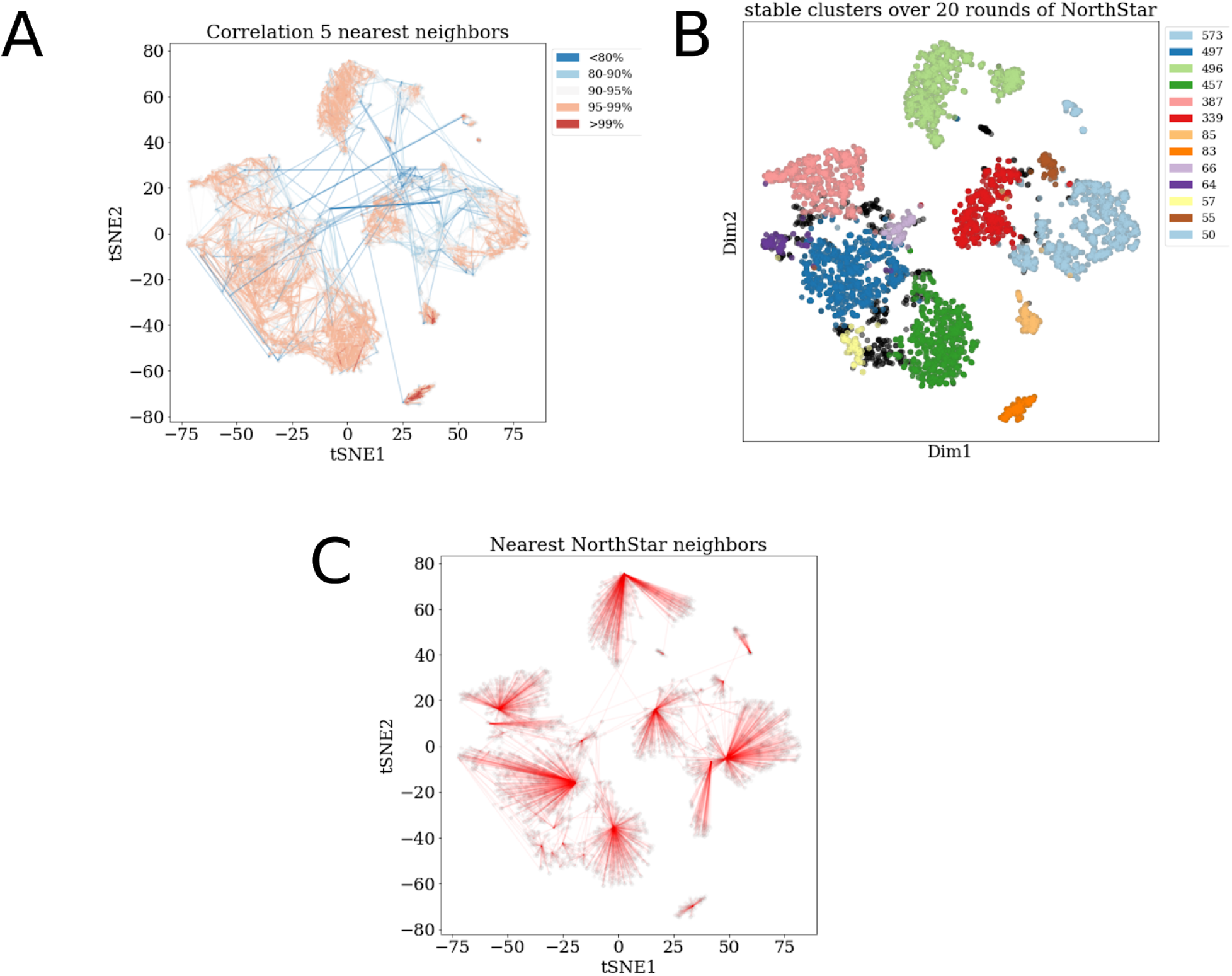
Distance matrix characteristics and cluster stability using northstar. **(A)** Edges of 5 highest correlating cells drawn for each cell, colored by correlation (= 1 - distance). This correlation matrix of cells and atlas class ‘cells’ was used to perform weighted PCA and render t-SNE as shown in Figure 2. Correlation is the pairwise correlation of each cell using the union of top 20 overdispersed genes for each atlas class and the top 400 overdispersed genes of the new glioblastoma dataset. **(B)**: Top 11 largest clusters of cells (colors and number of cells) that always grouped together over 20 rounds of re-initialization northstar (atlas cell weight=60, number of PCs=20, resolution parameter=0.0012. threshold neighborhood=.8, with self-edging). Cells forming smaller consistent clusters or that are grouped varyingly shown in black. **(C)**: Single nearest neighbor of each cell, based on averaged class-based correlation matrix taken over 20 rounds of re-initialization of northstar.

**Supplementary Figure 3.**
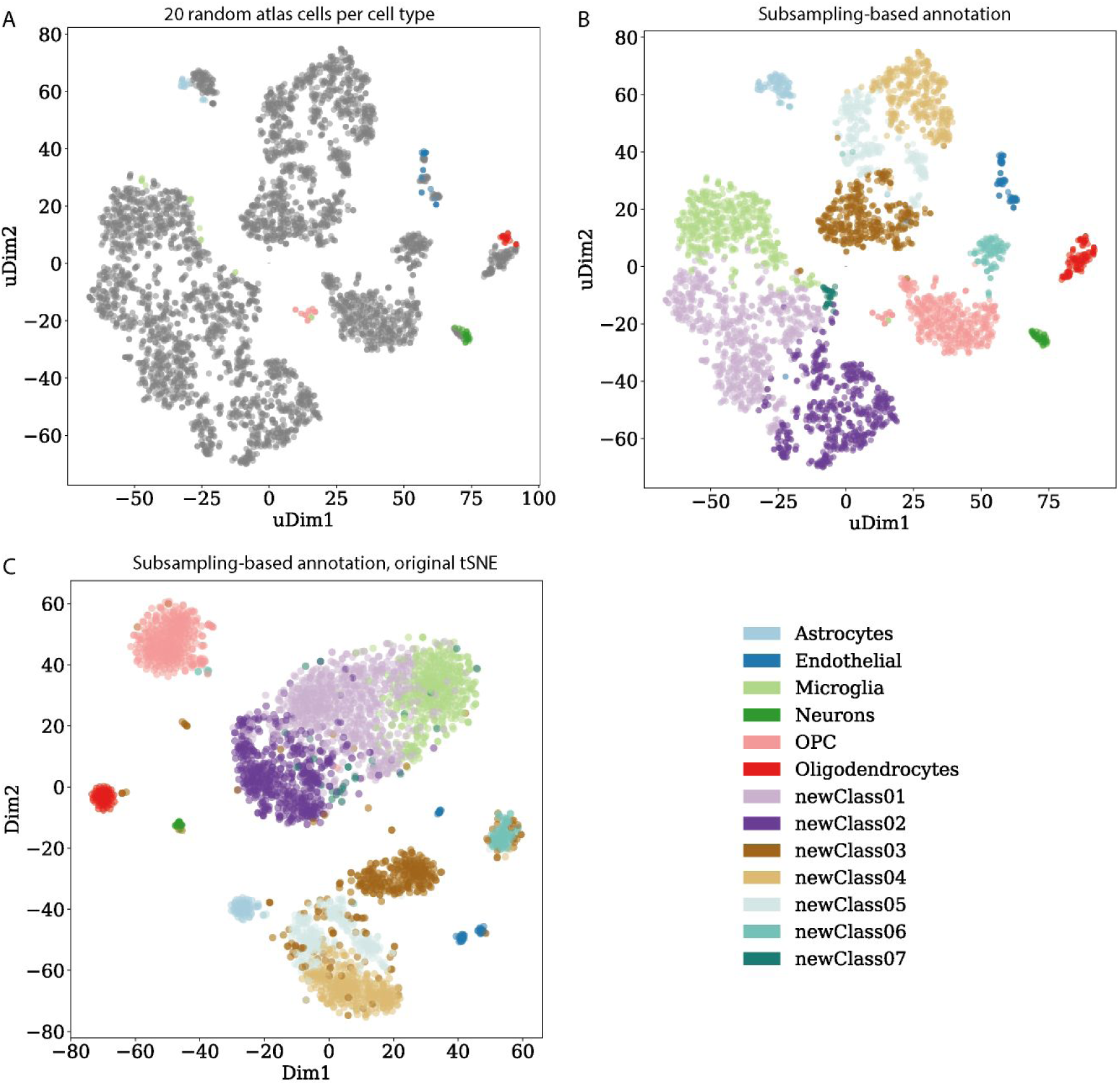
Annotation of the GBM dataset by subsampling atlas cells. Notice that the microglia annotation is expanded to include around a third of the cells that were originally annotated more generically as “immune cells”, in agreement with the result found by averaging within cell types.

**Supplementary Figure 4.**
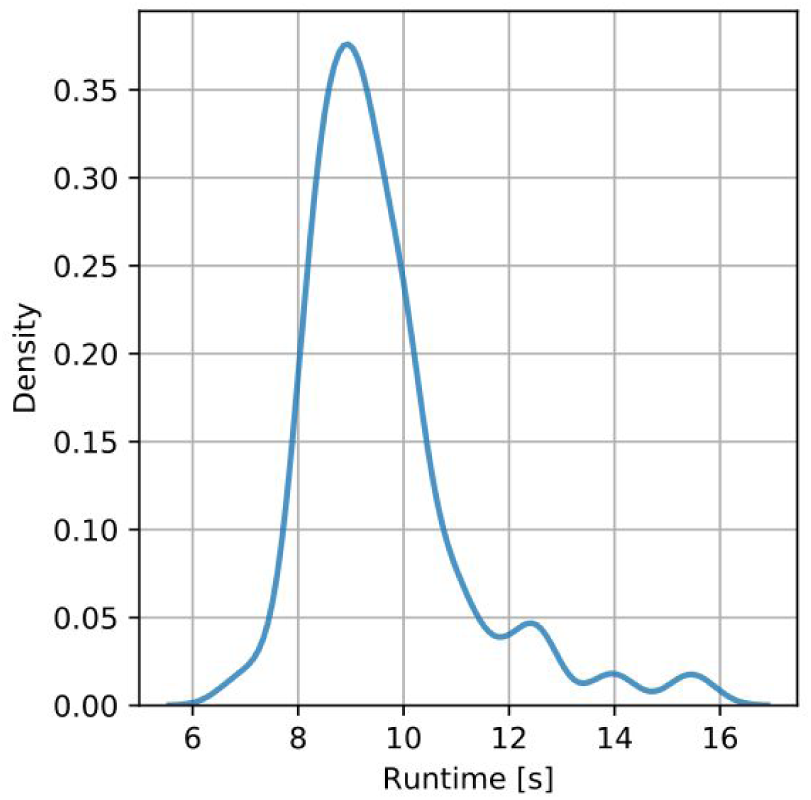
Runtime performance. Distribution of northstar runtimes for the melanoma dataset (>4,000 cells) in runs with accuracy >=80% (including with default parameters).

**Supplementary Figure 5.**
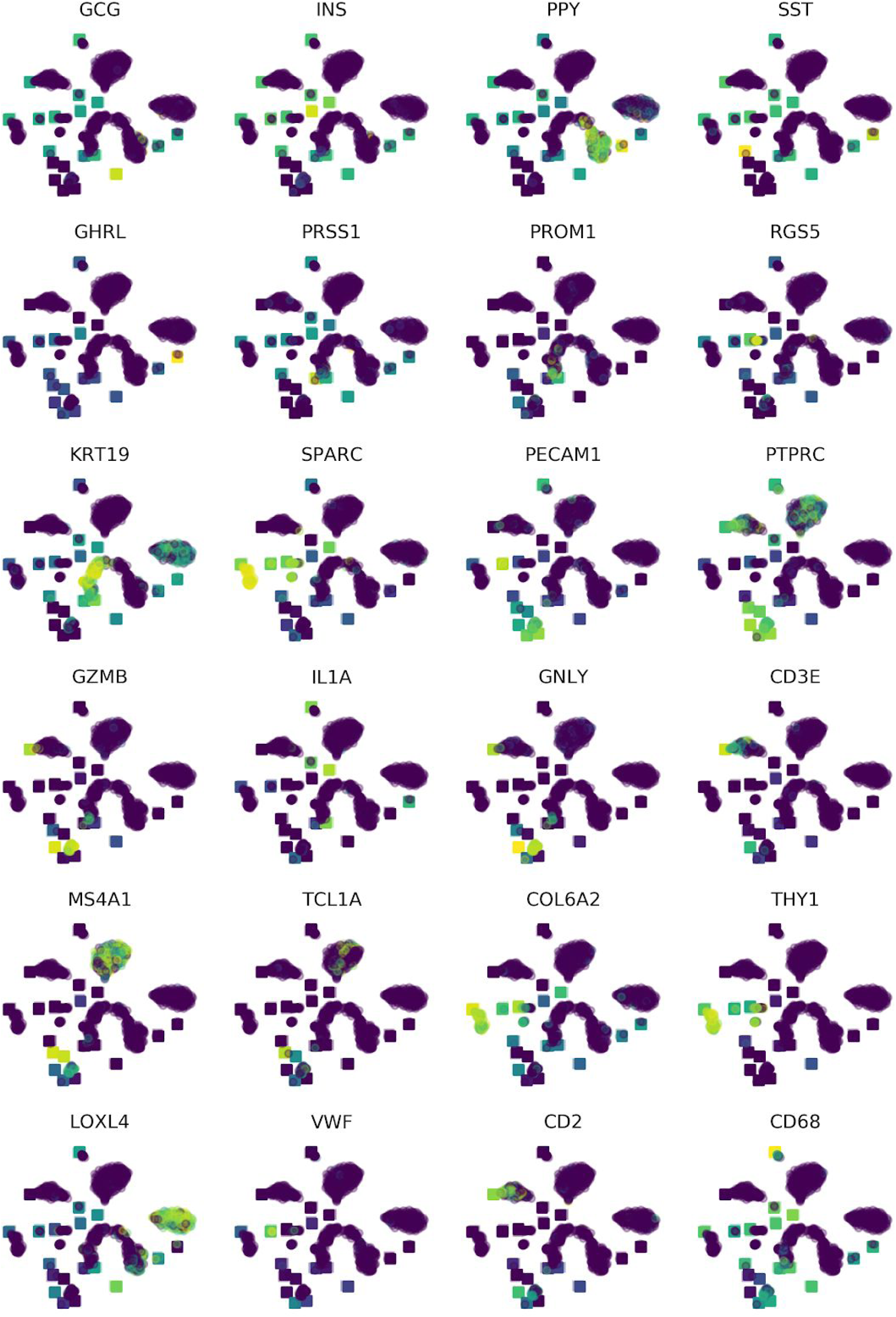
Key marker genes for different pancreatic and blood cell types. projected onto the t-SNE of Figure 3A, colored by their expression level (low: blue, intermediate: green, high: yellow). As in Figure 3A, squares indicate atlas cell types, circles new cells from pancreatic tumors.

### Supplementary Text 1: Detailed description of the northstar algorithm

#### Merge cell atlas and new single cell dataset

northstar starts with a gene expression matrix M with L rows (genes) and N columns (cells). The first N_a_ columns represent the cell atlas. Since the atlas is already annotated, the gene expression of each cell can be approximated by the average of its cell type. Therefore, each of these columns contains the mean gene expression within a cell type from the atlas. In addition to these averages, the number of atlas cells within each cell type is taken as input as a vector, which we call S. The last N_n_ = N - N_a_ columns of M represent single cells from the new dataset (“new cells”), which need to be annotated. For these columns, no equivalent of S is needed since each column describes only one cell. The input data for northstar is also depicted in Figure 1A. The steps below are illustrated in Figure 1B.

#### Feature selection

The first step of northstar is to select features (genes) to calculate the neighborhood graph. Briefly, feature selection is necessary because noisy points in a high-dimensional space are, loosely speaking, almost equidistant from one another (“curse of dimensionality”): the selection of 300-1000 most relevant genes drastically reduces noise while retaining enough information to calculate neighborhoods. It is empirically observed that excluding key features from the selection might cause overclustering, while erring on the side of too many features has a less dramatic impact on the results.

A common and simple way to select features is to take the most overdispersed genes, i.e. genes with a large variance-to-mean ratio across all cells. northstar by default takes this approach for the sake of simplicity, however it includes two modifications. First, it only computes overdispersion within the new dataset, since the atlas might be much larger and might otherwise dominate the feature selection, obfuscating de novo cell type discovery. Second, it explicitly includes the most discriminating genes for each cell type in the atlas, to ensure that new cells can be positioned correctly within the atlas if they do belong to any of the known cell types. To maximize customizability, the user can select features before northstar if she prefers to use different criteria.

#### Weighted PCA

Principal component analysis (PCA) is a commonly used technique across scientific fields and is used in single cell transcriptomics to further mitigate the effect of noise in high-dimensional spaces beyond feature selection. northstar performs PCA on the feature-selected data using weights to balance the influence of the atlas and the new dataset on the annotation result.

Note that most descriptions of PCA consider the rows as observations and columns as variables, whereas we consider a gene expression matrix M that has the cells as columns and the genes as rows, to be more consistent with the single cell transcriptomics literature (see above). In any case, the same operations can be trivially applied to the transposed matrix and lead to the same result.

First, the weights are normalized into fraction of the total weight via division by the total number of cells in both atlas and the new dataset. The resulting vector of weights is then written in a diagonal matrix W of size N x N. We call this matrix the normalized weight matrix.

Then, M is linearly shifted to have zero weighted mean across all cells. It is then normalized by dividing by the square root of the weighted variance across all cells.

The weighted covariance matrix of M is calculated as Cov_w_ M = M W M^T^. This matrix has dimensions L x L and is real and symmetric, so its canonical linear map has L real eigenvalues and L real eigenvectors. Let U be the matrix with the eigenvectors as columns.

The first P eigenvalues and eigenvectors are computed by standard linear algebraic techniques. Although all L components can be calculated in principle, it is in practice sufficient for the sake of subsequent steps to trim this operation to P ~ 20 because the spectrum of eigenvalues decays sharply after the first few elements.

The P eigenvectors are the top gene loadings of the weighted PCA. To find the cell loadings or principal components (PCs), we exploit the well known connection between PCA and singular value decomposition (SVD): the eigenvectors of Cov_w_ M are also right singular vectors of the data matrix M, while the PCs are left singular vectors. The matrix V with the PCs as columns can therefore be computed by matrix multiplication:

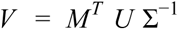

where Σ is the smi-diagonal matrix with the singular values (i.e. the square roots of the eigenvalues of the covariance matrix) as the first diagonal entries and zero elsewhere and Σ−^1^ indicates the reciprocal of all nonzero singular values and zeros elsewhere. In theory, it is possible that this inversion cannot be computed if the covariance matrix is degenerate (i.e. it has rank < L). However, such pathological cases are unlikely to appear in real datasets and are not considered further here.

Note that the linearity of PCA explains why it is possible to calculate an approximate result without having to load all atlas data: since the atlas cells are approximated with their cell type average, having a cell type with a weight of 2 is exactly the same as duplicating that column into the data matrix and having both copies have a weight of 1.

#### Construction of the k nearest neighbors graph

To construct the k nearest neighbors (knn) graph, a distance matrix from the new cells and all cells (including atlas and new ones) is first computed. The software package lets the user choose the distance metric and defaults onto Pearson correlation, a commonly used metric in single cell transcriptomics. The distance matrix has dimensions N_n_ x N.

Then, for each new cell (row in the distance matrix) the k elements at closest distance excluding self are identified. Notice that there is no need to sort the distance vector fully for this. The algorithm then scan these candidate neighbors in order of increasing distance. If a candidate belongs to the new dataset, it is added to the neighbors list of this cell. If a candidate belongs to the atlas, however, it is a representative of a larger cell type which is bona fide tightly clustered around it. Hence, not only one neighbor is added but rather a number equal to the size of the cell type in the atlas. The total number of neighbors is then trimmed to k.

In addition to the neighbors from the new data into the atlas, adding a few edges that are computed outwards from the atlas helps to reduce batch effects (simlarly to mutual nearest neighbors schemes). We usually compute 5 neighbors in the new dataset for each atlas average and add this small amount of edges to the similarity graph.

It is possible that a cell (or atlas average) has no close neighbors or just fewer than k. To consider this situation, a maximal distance threshold is used to compute the neighbor candidates. The default for correlation distance is 0.8: cells with a Pearson r < 0.2 with the current cell of interest are never treated as neighbors.

Once the nearest neighbors for every new cells are found, an undirected graph is constructed by symmetrization of all edges. Edges between new cells and atlas cell types can be present multiple times in the neighbors lists and are weighted accordingly. This increased edge weight ensures that if a cell is really close to a known cell type it will rapidly be absorbed into that cluster during the Leiden algorithm below. No direct edges are set between atlas cell types because they are annotated already and are not allowed to change cluster membership during the modified Leiden algorithm.

#### Leiden clustering with fixed nodes

The joint dataset is now represented as a neighborhood graph and can be clustered using standard graph-based methods. However, the nodes belonging to atlas cell types are already annotated and should not be allowed to change membership lest a full reannotation of the atlas is required.

northstar solves this issue by modifying the Leiden algorithm for community detection in large graphs to allow for a number of nodes to be “fixed” into their initial membership. The Leiden algorithm itself has been proven effective in unsupervised clustering of single cell transcriptomic data, scales well with graphs with millions of nodes, and provides mathematical connectivity guarantees that lend trust to the resulting annotations.

Briefly, Leiden performs two kinds of operations on nodes, namely “move” and “merge”. Moreover, it recursively collapses the initial graph into simplified aggregated graphs in which each cluster becomes a single node: the “move” and “merge” steps are then repeated in the aggregated graph. northstar changes both the “move/merge” steps and the aggregation step.

First, whenever a queue of nodes is constructed for consideration in terms of a move/merge, nodes that are marked as fixed are just never considered. This prevents them from switching membership to another community. Second, whenever aggregated graphs are constructed, fixed nodes are collapsed only if they belong to the same community from the beginning; otherwise they are never collapsed.

The output of northstar is a list of cluster memberships for each new cell. Numbers 0 to (Na - 1) represent extant cell types in the atlas (corresponding to the first Na columns of M) and higher numbers indicate new inferred cell types in the new dataset.

### Supplementary Text 2: Extended methods

#### Analysis of the GBM dataset based on the human brain cell atlas by Darmanis et al

First, we deleted any information on the cell type annotations on the cells from glioblastoma patients. The annotations for the cells from the healthy brain atlas, excluding fetal cells, were kept as a training set and the gene expression of the atlas was approximated by averaging within each cell type. We then performed feature selection by taking a constant number of overdispersed genes (adjustable, here 20) for each atlas cell type as well as the top overdispersed genes of the new glioblastoma cells (typically the top 500 [currently shown: top 400]) to generate a feature-selected matrix. This matrix has the dimensions of at most 5*20+400 genes by 5 cell types + 3589 new cells, though it likely has fewer genes due to genes shared between cell classes or between the atlas and the new dataset. This matrix was used to instantiate and run northstar, and for visualization purposes we created a distance matrix by calculating the pairwise correlation between cells and atlas classes, followed by a weighted PCA that gives the atlas cells the weight of 60 single cells and using the top 20 PCs to generate a 2D rendering of cell similarity using t-SNE.

Projecting the original glioblastoma annotation back onto the t-SNE, we observe that our unbiased feature selection method roughly reproduces the previously reported cell clusters (Figure 2C, Supplementary Figure 1).

